# Engineered patient-derived organotypic bone model for multimodal characterization of donor-specific response to therapy

**DOI:** 10.1101/2025.04.17.649276

**Authors:** Anke de Leeuw, Chris Steffi, Gian Nutal Schädli, Amit Singh, Pei Jin Lim, Marianne Rohrbach, Cecilia Giunta, Matthias Rüger, Ralph Müller

## Abstract

Bone-forming therapies often fail in genetic skeletal disorders, highlighting critical gaps in mechanistic understanding and therapy evaluation. We developed 3D bioprinted organotypic bone models using primary cells from a patient with FKBP10-related osteogenesis imperfecta (OI) and from metabolically healthy controls obtained via femoral osteotomy (FO). Cyclic mechanical loading and reseeding generated patient-derived bone-like tissue for structural, molecular, and transcriptomic characterization of the engineered donor-specific tissue material. Dickkopf-1 antibody (DKK1Ab) was then administered as a therapeutic perturbation. OI constructs showed a bidirectional interferon-stimulated gene (ISG) signature and hypermineralization with structural fragility, hallmark features of OI. Unlike FO constructs, DKK1Ab administration in OI resulted in a limited transcriptional response marked by ISG downregulation and increased MKI67 expression. Therapeutic perturbation with DKK1Ab increased early procollagen I secretion, and was associated with lower fracture scores, although the within-OI difference was not statistically significant. This proof-of-concept demonstrates multimodal donor-specific characterization of engineered patient-derived bone models following DKK1Ab perturbation.

## 1 Introduction

Clinical translation of bone-forming therapies for rare genetic skeletal disorders remains challenging because improvements in bone mineral density do not translate into improved bone quality or fracture resistance. Recent randomized clinical trials in osteogenesis imperfecta, including TOPaZ^1^ and the pivotal ORBIT/COSMIC^2^ studies, further highlight this translational gap by demonstrating improved bone mineral density without consistent reductions in fracture risk. Mechanically stimulated organotypic bone models generated from patient-derived cells produce donor-specific bone-like tissue material, enabling structural, mechanical, molecular, and transcriptomic characterization following therapeutic perturbation.

Osteogenesis imperfecta (OI) represents an important proof-of-concept disease in which this challenge is particularly evident because of its marked phenotypic and genetic heterogeneity. Children with OI exhibit varying degrees of bone deformities and functional disability and experience a lifelong predisposition to unpredictable fractures. Despite decades of research, recent clinical trials have failed in providing protection against skeletal fragility despite preclinical models showing improvements in bone mineral density^1,2^. Patient-derived biofabricated organotypic *in vitro* bone models^3^ now offer unprecedented opportunities to decode these hidden mechanisms, because they enable comprehensive characterization of the mineralized extracellular matrix generated by patient-derived cells through complementary modalities including micro-computed tomography (micro-CT), histology, immunohistochemistry, structural, mechanical, molecular, and transcriptomic analysis^4^. This shifts the focus from developing bioinks, cell-laden hydrogels, and biomaterial-derived 3D in vitro cultures toward characterizing the functional properties of the patient-derived bone-like material generated within these systems.

Despite its critical importance for bone homeostasis and remodelling, mechanical stimulation has not yet been integrated into 3D *in vitro* cell cultures of OI. While recent work has demonstrated that mesenchymal stem cells from OI patients retain sensitivity to short-term shear stress in two-dimensional systems,^5^ such approaches do not recapitulate the dynamic, mechanically loaded bone microenvironment required to generate and functionally characterize mineralized extracellular matrix over time. Cyclic mechanical loading drives osteoblast-to-osteocyte differentiation,^3,6^ regulates mineralization patterns, and shapes bone microarchitecture^7^— all crucial factors in OI pathophysiology. Previous *in vitro* models have neglected this essential component,^8–11^ limiting their ability to examine loading-dependent matrix formation and responses to experimental perturbations.^12^

Currently, no FDA-approved treatments exist specifically for OI. Symptoms are managed with bisphosphonates, physical therapy, and surgical intervention. Recently, targeted anabolic protein therapies, such as sclerostin and Dickkopf-1 antibodies (DKK1Ab), have emerged as promising candidates to enhance bone formation by modulating the Wnt signalling pathway.^13–15^ Preclinical studies have demonstrated that DKK1Ab antibody therapy increases bone mass, mineral density, and improves bone structure in rodent models.^16,17^ Notably, children with OI exhibit significantly elevated serum DKK1 levels, both in untreated patients and those receiving bisphosphonate therapy.^15,18,19^ When healthy mesenchymal stem cells were cultured with serum from OI patients, osteoblast differentiation was inhibited; however, the addition of DKK1Ab antibodies reversed this effect, increasing the expression of RUNX2 and collagen I mRNA.^15,19^ Despite the potential of DKK1 antibodies to strengthen bone and reduce fractures, their effects on patient-derived OI cells remain unexplored in *in vitro* models. The molecular mechanisms underlying variable therapeutic responses in skeletal disorders remain poorly understood. Interferon-stimulated genes (ISGs), which normally function in antiviral defence, are also expressed in osteoblasts, where chronic activation of type I interferon responses suppresses osteogenesis. The contribution of individual ISGs differs. ^20^ISG STAT1 suppresses osteoblast differentiation by sequestering the master transcription factor RUNX2.^21^ ISG15 acts in the opposite direction, since its knockdown reduces alkaline phosphatase activity, osteogenic gene expression, and matrix mineralization in primary osteoblasts.^22^ An interferon signature in bone therefore requires resolution at the level of individual genes rather than interpretation as a single inhibitory program. Elevated interferon signatures, including IFI27, ISG15, and STAT1 have been documented in the blood of OI patients,^23^ suggesting that chronic immune activation may compound the skeletal defects caused by primary genetic mutations. However, the therapeutic potential of targeting interferon pathways in bone diseases remains largely unexplored.

A gelatine/alginate/graphene oxide biofabrication platform has been previously established and optimized over several works^3,6,24–26^. The bioink was optimized on rheology, printability, and swelling across a series of earlier studies. An amount of 0.8% alginate was identified as the concentration that preserved print fidelity and ionic crosslinking while giving the highest cell viability and the most dendritic cellular morphology, by varying the concentration from 0.8 to 2.3%.^26^ Comparing scaffold stiffness, the softer 0.8% matrix (0.66 kPa) produced higher ALP activity, stronger osteogenic differentiation, and nearly twice the mineralized tissue by day 42 relative to a stiffer 1.8% matrix (5.4 kPa), establishing the low-stiffness formulation as optimal for bone tissue engineering applications.^25^ Introducing graphene oxide across a 0.5 to 2 mg/mL series, we found that the 1 mg/mL graphene-oxide composite maximized bioprintability, scaffold fidelity, compressive modulus, and osteogenic gene expression, improving reproducibility and osteogenic properties.^24^ Because graphene oxide concentration was varied independently of the fixed alginate/gelatin composition, rheological testing directly attributed the increases in ink viscosity, shear-thinning, and shear recovery to graphene oxide.^24^ Optimizing the mechanical stimulation protocol, it was found that long-term cyclic loading applied from day 1 maximized mineral density, construct stiffness, and osteoblast differentiation and drove collagen-I maturation, osteocyte differentiation, and lacunar-canalicular network formation, establishing a human mesenchymal stem cell-derived osteocyte bone organoid.^6^ Translating the platform to primary patient-derived osteoblasts, a physiological printing density of 10 × 10 cells/mL accelerated mineral formation and increased construct stiffness roughly ten-fold over the culture period, compared with about two-fold at lower density, optimizing the cell density for creating organotypic patient-derived bone models.^3^

Here, the platform first matures cell-laden hydrogel constructs using patient-derived osteoblasts under cyclic mechanical loading within four weeks,^3,6^ followed by osteoblast reseeding. Primary osteoblasts are subsequently cultured on the mineralized constructs, enabling continued deposition and modelling of patient-derived mineralized extracellular matrix that can be structurally, mechanically, molecularly, and transcriptomically characterized.

We focused our study on recessive FKBP10-related osteogenesis imperfecta as a representative of a particularly under-studied and neglected subset of OI with a high unmet medical need. Murine models consistently fail to recapitulate human OI phenotypes, particularly in subtypes like FKBP10-related OI where global FKBP10 knockout mice are perinatally lethal and conditional osteoblast-specific knockouts, while viable, do not fully recapitulate the skeletal defects observed in severe OI.^27–29^ Patient-derived cells are therefore essential to study disease-relevant phenotypes. A recent cross-species analysis integrating single-cell transcriptomics of the endosteal compartment with an extended genome-wide association study of estimated bone mineral density reported that known OI and bone fragility genes are overrepresented in pre-osteoblasts and mature osteoblasts, supporting the osteoblast lineage as the relevant cellular context for these disorders.^30^ Insights gained from recessive OI may also complement studies of dominant collagen I-related forms, contributing to a broader understanding of osteogenesis imperfecta. Using patient-derived osteoblasts from *FKBP10*-related OI and metabolically healthy controls from femur osteotomy, we evaluated whether our mechanically stimulated biofabrication platform could generate donor-specific bone-like material suitable for multimodal structural, mechanical, molecular, and transcriptomic characterization following DKK1Ab perturbation. Rather than establishing a generalized disease model, this proof-of-concept study investigates whether our mechanically stimulated biofabrication platform can reproducibly generate donor-specific bone-like material whose structural, mechanical, molecular, and transcriptomic properties provide functional surrogate readouts of therapeutic response. This proof-of-concept study focuses on the generation of donor-specific bone-like material and on its structural, molecular, and transcriptomic characterization following DKK1Ab perturbation.

## 2 Results

### 2.1 Biofabrication and mechanical loading of patient-derived 3D bone models for treatment evaluation

In this study, a mechanically stimulated 3D bioprinted organotypic bone model was developed to replicate bone tissue architecture and assess patient-specific responses to osteoanabolic treatment. Primary osteoblasts isolated from bone explants from an FO and an OI patient with *FKBP10*-related OI (OI) were expanded and encapsulated in an osteogenic bioink for extrusion-based 3D bioprinting (Fig. 1A). To promote mineralization and osteogenic differentiation, bioprinted constructs were subjected to a 4-week mechanically loaded preculture period with uniaxial cyclic compressive loading. At the end of preculture, mineralized constructs were reseeded with fresh primary osteoblasts from the same patient, allowing newly introduced cells to interact with mineralized surfaces, mimicking osteoblast-osteocyte communication *in vivo*. Following reseeding, constructs were cultured for an additional 4 weeks under adaptive mechanical loading, in the presence of either 1 µg/mL Dickkopf-1 antibody (DKK1Ab), an inhibitor of the Wnt signalling antagonist DKK1, or 1 µg/mL goat IgG isotype control (IgG) to account for non-specific antibody effects (Fig. 1B).

**Fig. 1.**
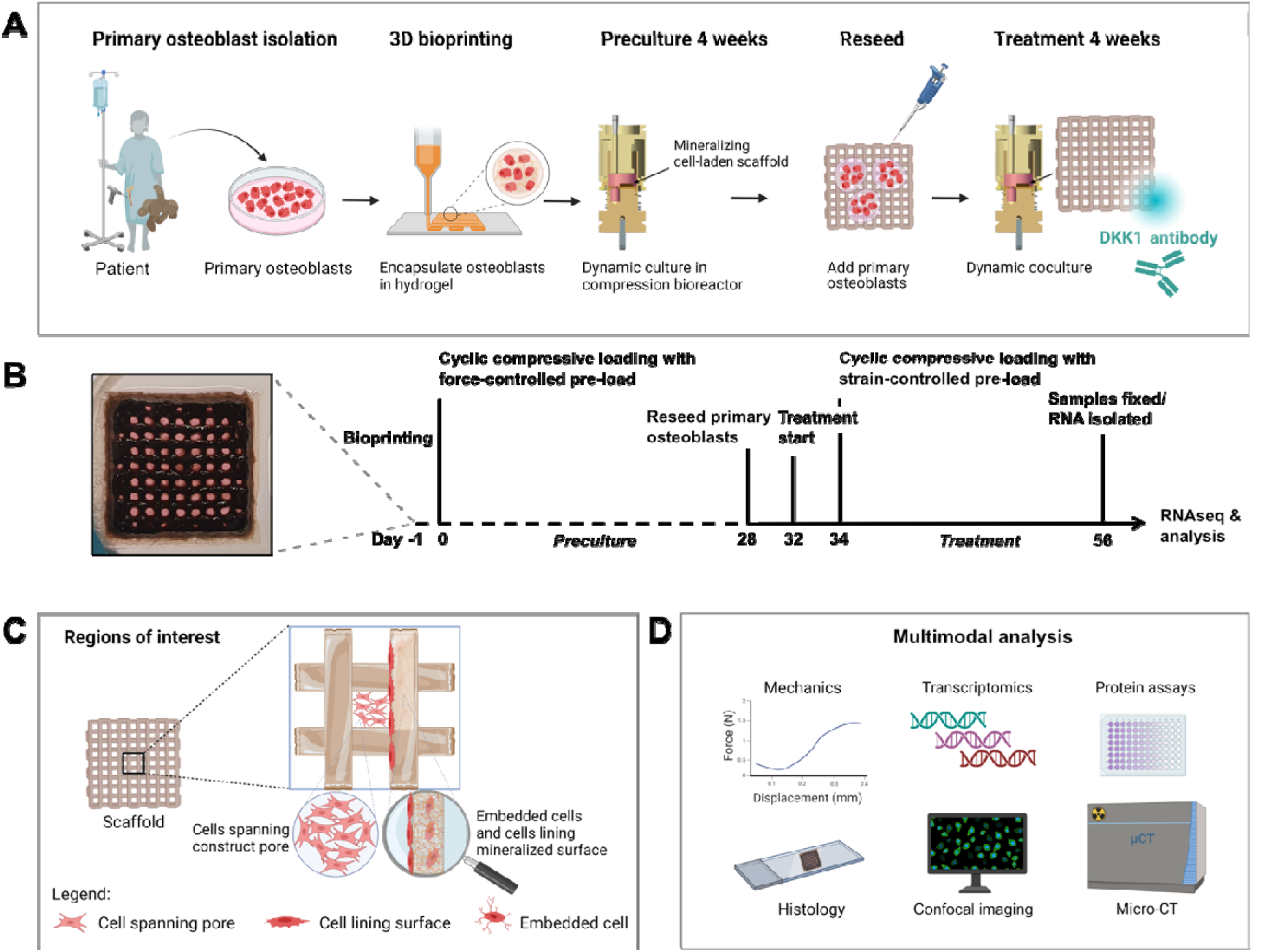
Organotypic bone model study design. (A) Primary osteoblasts from a femoral osteotomy (FO) and an osteogenesis imperfecta (OI) patient’s bone explants were isolated, expanded and encapsulated in osteogenic bioinks for extrusion 3D bioprinting. Cell-laden constructs were matured during 4-week preculture under uniaxial cyclic compressive loading and reseeded by pipetting fresh primary osteoblasts from the same patient on mineralized surfaces. Samples were then cultured in the presence of 1 µg/mL Dickkopf-1 antibody (DKK1Ab) or 1 µg/mL goat immunoglobulin G (IgG) isotype control under compressive loading for a further 4 weeks. The schematic was generated with BioRender. (B) Experimental timeline showing bioprinted organotypic bone model, start of 4-week mechanically loaded preculture period with fixed loading (0.07 N contact position preload, and cyclic compressive loading 1 % strain 5 Hz 300 s) followed by reseeding fresh osteoblasts, start of treatment and adaptive mechanical loading (0.05 N contact position + 10 % strain preload, and cyclic compressive loading 1 % strain 5 Hz 300s) to avoid overloading the scaffolds. (C) Schematic showing regions of interest in 3D bioprinted organotypic bone models, including mineralized surfaces of bioprinted struts and macroscale scaffold pore regions. (D) Multimodal analysis of organotypic bone models including mechanics, histology, transcriptomics, confocal imaging, protein assays and time-lapsed micro-CT scans.

The organotypic bone model was designed to replicate native bone modelling, with bioprinted cell-laden filaments undergoing mineralization and subsequently recruiting fresh osteoblasts to mineralized surfaces upon reseeding. These newly introduced osteoblasts spanned the macroscale scaffold pores or lined the mineralized struts, contributing to extracellular matrix deposition and tissue maturation (Fig. 1C). To evaluate the impact of combined mechanical loading and DKK1Ab treatment on osteogenesis, a multimodal analysis workflow was applied, integrating time-lapsed micro-computed tomography (micro-CT), fracture analysis, histological assessments, confocal imaging, transcriptomic profiling, and protein quantification (Fig. 1D).

### 2.2 OI organotypic bone models exhibit increased mineralization and loss of construct fidelity

Investigating mineralization dynamics in patients through repeated CT scans is clinically not feasible due to radiation exposure and logistical constraints. Organotypic bone models offer a patient-specific alternative, allowing longitudinal monitoring of mineralization through weekly time-lapsed micro-CT imaging (Fig. 2, Supplementary Videos 1-4). 3D reconstructions revealed fractures and a loss of fidelity in OI organotypic bone models (Fig. 2A). Loss of fidelity and deformations in the bioprinted lattice structure became apparent after day 35 in OI samples. Visually, treatment with Dickkopf-1 antibody did not appear to reduce the fractures in the scaffold struts but did appear to reduce the severity of deformations in OI organotypic bone models (Supplementary Videos 3-4). Small, mineralized structures appeared in the macroscale pores of organotypic bone models after day 35, particularly evident in the intact pores of FO samples (Supplementary Videos 5-6). In OI constructs, mineral volume at day 21 and 28 (Fig. 2B) was increased (day 7-14) due to accelerated early mineral formation compared to FO samples (Fig. 2C). However, between day 28 and 35 mineral formation in OI dropped significantly, coinciding with loss of fidelity, leading to a plateau in mineral volume in both treated and control groups. Mineral density was significantly higher in OI groups compared to FO groups throughout the duration of the study (Fig. 2D). Mechanical testing showed no significant differences in stiffness among groups at endpoint (Fig. S1A). In FO models, stiffness correlated positively with mineral density, consistent with previous findings.^6^ In OI constructs, stiffness and mineral density were uncorrelated.

**Fig. 2.**
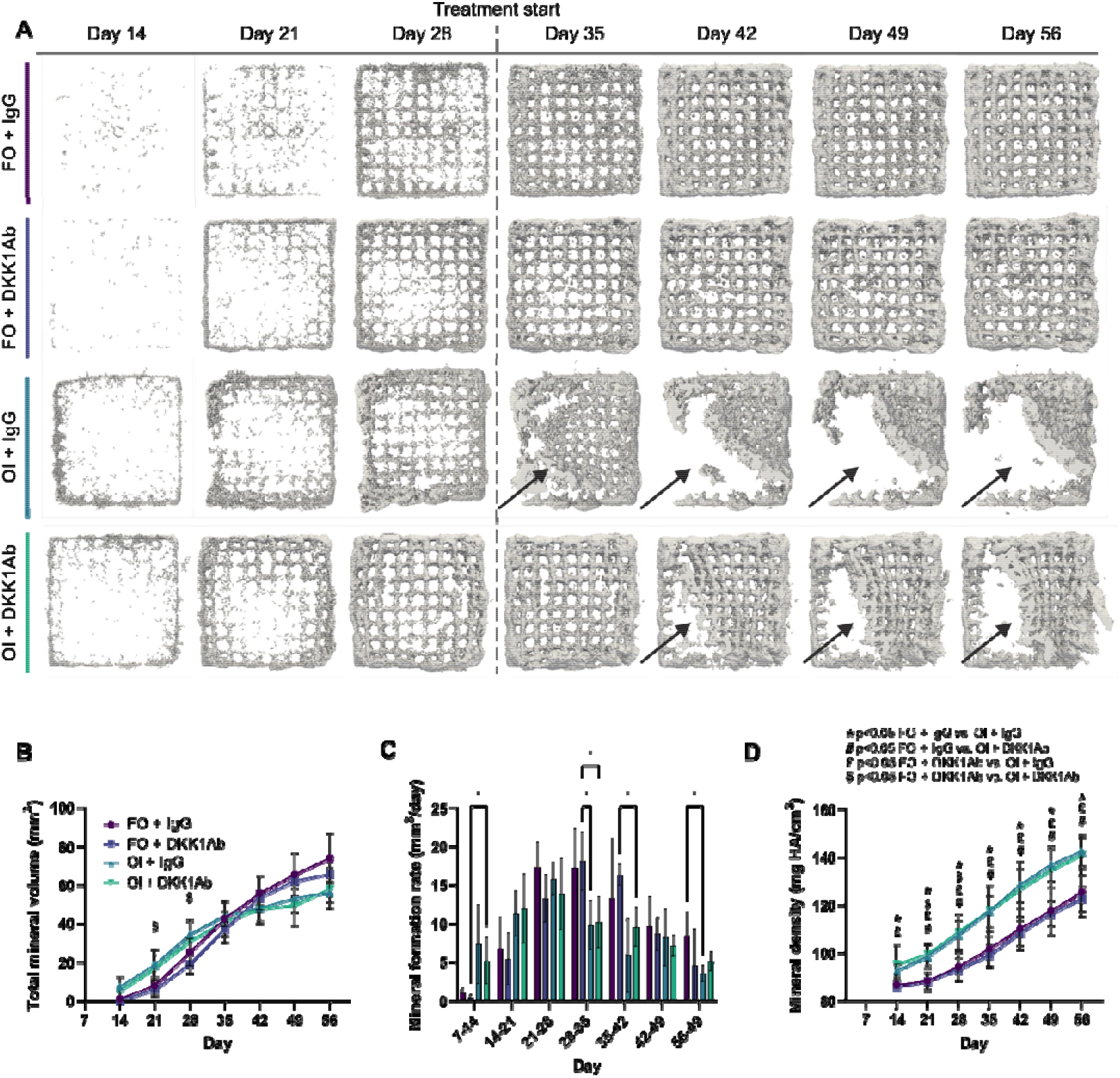
Longitudinal monitoring of mineralization and endpoint stiffness of organotypic bone models created with primary cells from a femoral osteotomy (FO) and an osteogenesis imperfecta (OI) patient. Treatment with isotype immunoglobulin (IgG) control and Dickkopf-1 antibody (DKK1Ab) commenced on day 32. (A) Time-lapsed 3D reconstructions of representative organotypic bone models, arrows show regions affected by fractures and deformity. (B) Total mineral volume, (C) Mineral formation rate (mineral volume changes over time) and (D) Mineral density. Data are shown as mean ± standard deviation (n ≥ 5). Statistical analysis was performed using two-way ANOVA followed by Tukey’s multiple comparisons test. Symbols indicate statistical significance p<0.05; * was used to compare mineral density between patients, i.e. FO + IgG with OI + IgG; # was used to compare FO + IgG with OI + DKK1Ab; £ was used to compare FO + DKK1Ab versus OI + IgG; $ was used to compare FO + DKK1Ab versus OI + DKK1Ab.

### 2.3 Biomimetic co-culture with distinct phenotypes between cells spanning pores and cells lining mineralized surface

To capture phenotypic differences in ECM organization, we performed histological (Fig. 3) and histochemical (Fig. S2) staining on patient-derived organotypic bone models cultured under cyclic loading for 8 weeks. Both FO and OI organotypic bone models were reseeded mid-culture to provide new osteoblasts for mineralized surfaces and pore regions. Histological overviews (Fig. 3A) identified regions of mineralized surfaces and pores within the same 3D model. H&E staining showed that cells on mineralized surfaces adopted flattened morphologies, while those spanning construct pores appeared more fibroblastic (Fig. 3B-E, H&E). This differential cell localization and morphology were accompanied by a different pattern of ECM secretion. Picrosirius red staining for investigating collagen deposition showed dense cell-secreted collagen deposition on mineralized surfaces and fibrillar, woven-like ECM spanning pores (Fig. 3B-E, Picrosirius Red). OI organotypic bone models (Fig. 3C) produced more collagen in pore regions compared to FO constructs (Fig. 3B). Alizarin Red S for calcium deposition showed homogeneous ECM staining in pores of FO constructs (Fig. 3B, Alizarin Red S) compared to the distinct mineral nodules in pores of OI samples (Fig. 3C, indicated by arrows). Immunohistochemical collagen type I antibody staining showed higher signal in OI organotypic bone models (Fig. S2). Treatment with DKK1Ab appeared to result in more dense mineralized collagenous ECM spanning pores (Fig. 3D-E, Fig. S2). Overall, these results indicate that this biomimetic co-culture method supports the formation of mineralized organotypic bone models mimicking features of woven bone and surface formation and reveals differences in OI and FO patient-derived models.

**Fig. 3.**
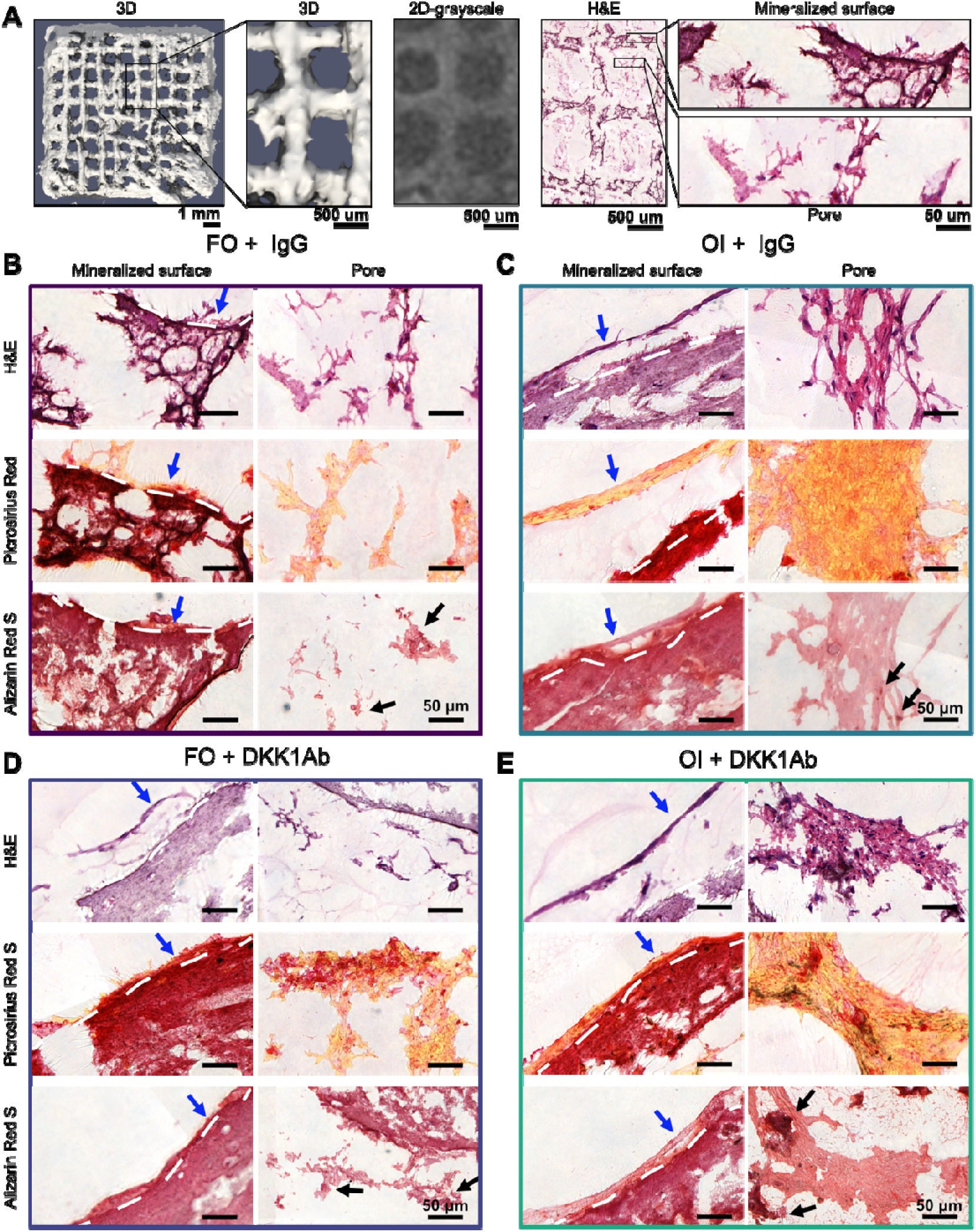
Extracellular matrix phenotypic characterization of FO control and OI patient organotypic bone models. After 8 weeks of compression loading, consecutive sections of FO and OI organotypic bone models were stained to visualize cell and extracellular matrix morphologies in the same region. (A) Overview where regions of interest (mineralized surface, pore) are located within the 3D construct. (B-E) Sequential histological sections of hydrogel interface and pore regions in FO (B,D) and OI (C,E) patient organotypic bone models treated with IgG control or DKK1Ab. Sections were stained with Haematoxylin and eosin (H&E) for showing morphology and localization of cells; with Picrosirius red for collagen (red-orange) on the construct surface and spanning across pores; with Alizarin Red S for mineralized cell-secreted collagenous extracellular matrix; mineral nodules are indicated by black arrows and bone lining-like cells on mineralized surface by blue arrows.

### 2.4 Organotypic bone models responses to DKK1Ab treatment

As fractures and deformities emerged in the OI organotypic models (Fig. 2A), we applied an image-processing method on time-lapsed micro-CT images to quantify volumetric changes over time (Fig. 4). Fig. 4A displays the changes of engineered tissue volume, categorizing volumes as Deformed (yellow) or Stable (red) relative to the day 28 baseline. FO organotypic bone models maintained remarkable integrity throughout the experiment, with both treatment conditions exhibiting minimal volumetric changes. In contrast, OI organotypic bone models demonstrated profound structural deformity.

**Fig. 4.**
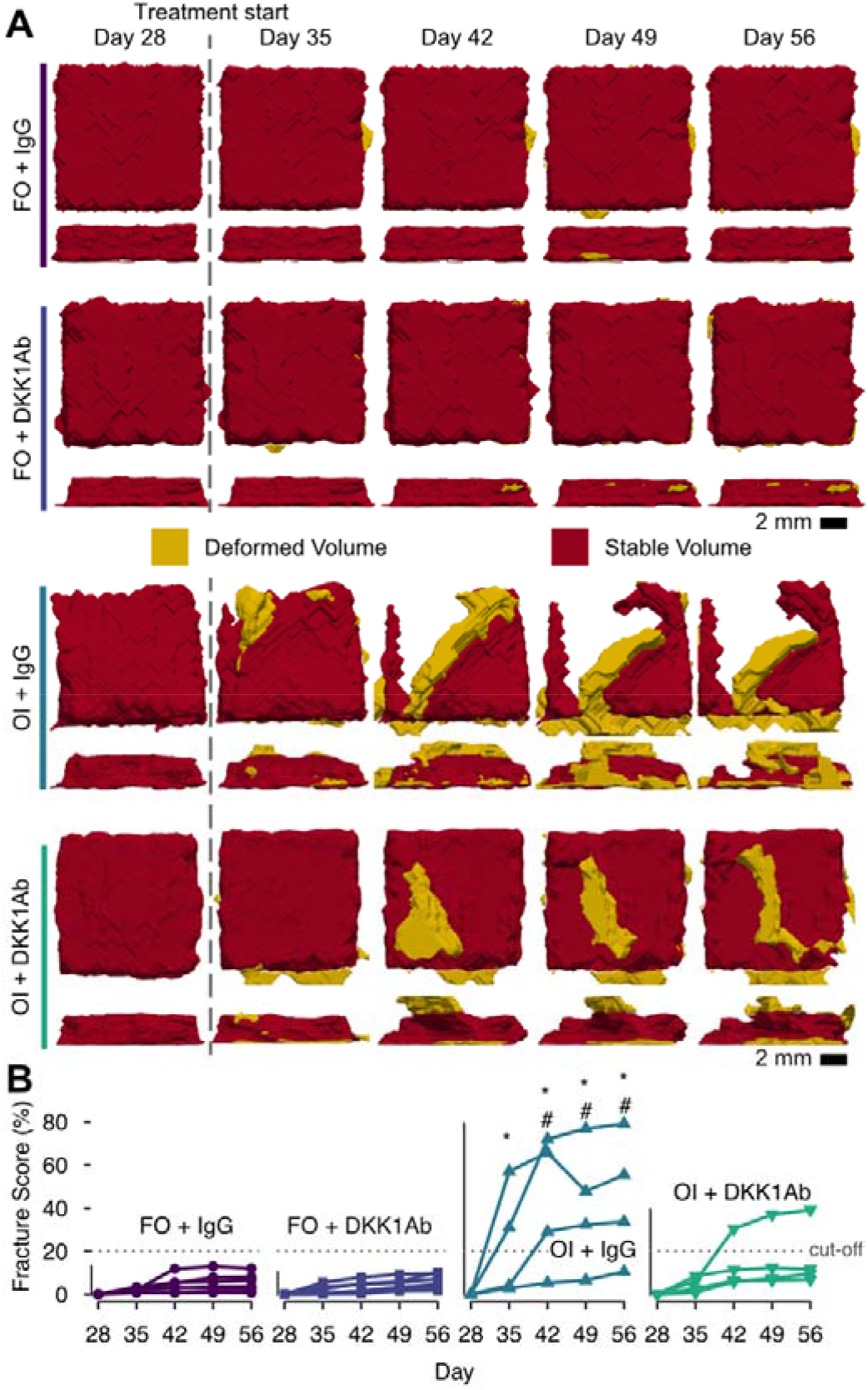
Fracture score analysis in FO and OI organotypic bone models treated with IgG or DKK1Ab. (A) Registered time-lapsed micro-CT images from day 28-56 for the four experimental conditions showing Deformed Volume (yellow), Stable Volume (red) relative to the day 28 reference. (B) Quantitative fracture score analysis for the four experimental conditions. FO + DKK1Ab and FO + IgG, n = 6 samples; OI + IgG and OI + DKK1Ab, n = 4 samples. Data represent individual samples. Statistical significance was assessed using nparLD with Benjamini-Hochberg post-hoc tests. Letters indicate significant differences (p < 0.05) as follows: *, different from FO + IgG; #, different from FO + DKK1Ab.

The resulting fracture score integrates volumetric deformation and trabecular strut thickness and spacing to capture progressive structural failure within each sample (Fig. 4B). FO models remained structurally stable, with fracture scores near baseline through the 4-week treatment period; median 2.7% for FO+IgG and 2.2% for FO+DKK1Ab. The OI+IgG models deteriorated starting from week 5 to 8 (median 30.0%, maximum 79.2%), whereas DKK1Ab mitigated deterioration and reduced the fracture score (median 6.7%, maximum 39.2%). Statistical analysis confirmed that fracture score progression was significantly higher in OI + IgG compared to FO+IgG (p = 0.043 at day 56) and FO+DKK1Ab (p = 0.034), while no difference between OI+DKK1Ab and any FO group was found at a pairwise timepoint level. Thus, DKK1Ab treatment attenuated the fracture scores in OI constructs toward FO levels, but did not eliminate structural fragility in the OI constructs.

Surface and pore mineralization dynamics inferred by reseeded cells could only be assessed in constructs that maintained structural integrity, therefore, this analysis was restricted to FO organotypic bone models (Fig. 5). Because DKK1Ab treatment was initiated four weeks into the experiment, when the bulk of the 3D bioprinted constructs had already mineralized, drug response might have been more pronounced in reseeded cells. We therefore applied an image-processing method that is more specific to surface specific mineralization and is also sensitive to mineralization induced by cells spanning pores (Fig. 3).

**Fig. 5.**
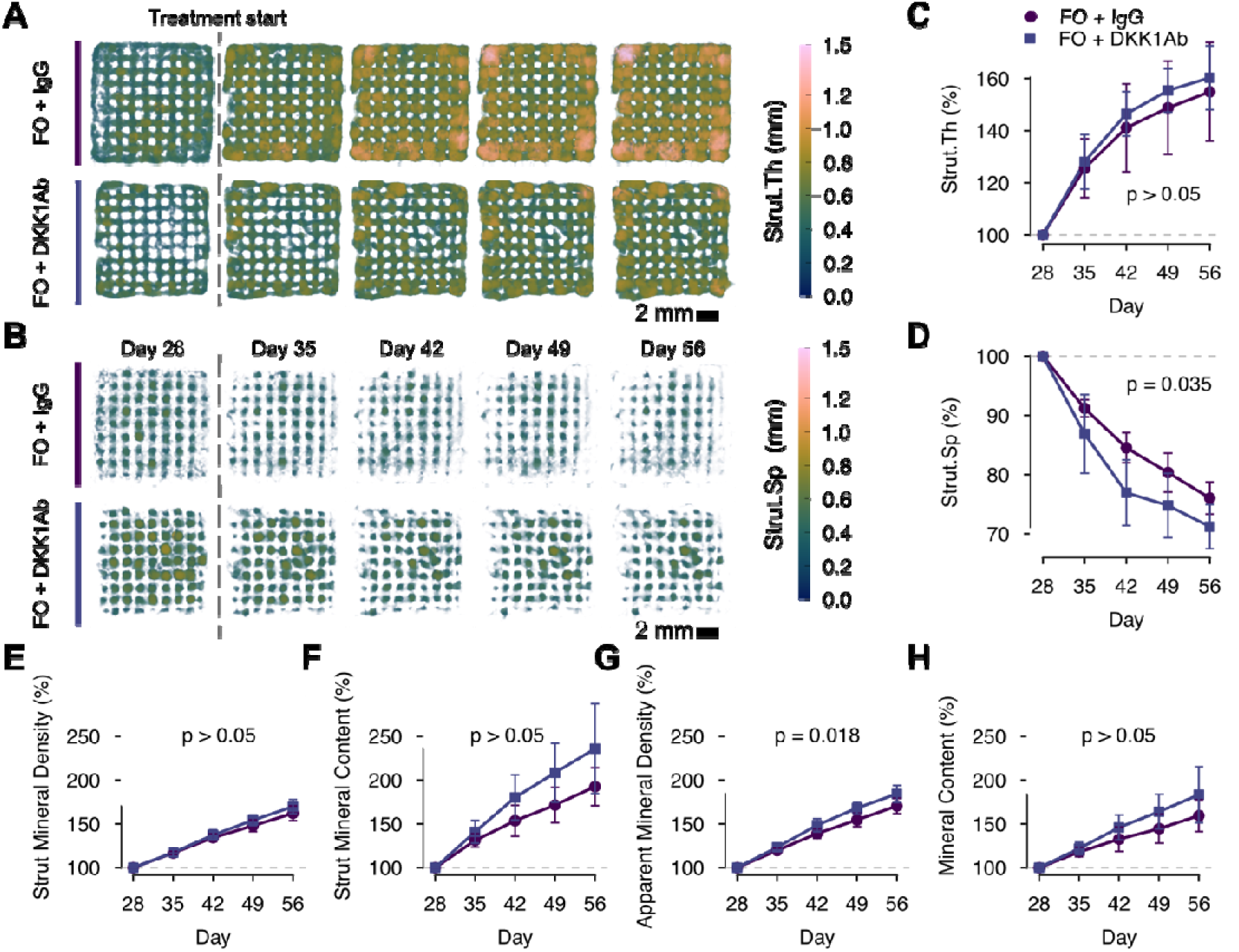
Surface mineralization analysis in FO organotypic bone models. (A) Registered time-lapsed micro-CT images from day 28–56 showing the evolution of strut thickness in FO + IgG (circles) and FO + DKK1Ab (squares) constructs. Colour maps represent local strut thickness (Strut.Th) and (B) corresponding visualization of strut spacing (Strut.Sp) progression. (C–D) Quantitative analysis of average strut thickness and spacing over time. (E,F) Strut mineral density and content quantified within the mask of the strut thickness and (G,H) apparent mineral density and mineral content quantified within the mask, representing the union of strut thickness and spacing. Data are mean ± SD for n = 6 per group. Statistical significance was assessed using repeated measures two-way ANOVA (E-H), and p-values < 0.05 report a significant difference between groups (C-H).

To further explore whether reseeded cells increase extracellular matrix deposition and mineralization with DKK1Ab treatment we separated mineralization effects within the pores from those within the bulk structure by creating a mask for the mineralized struts (Fig. 5A) and the strut spacing (Fig. 5B). The strut thickness increased significantly (Fig. 5C), shown by more yellow to pink colour (Fig. 5A). Although the FO+DKK1Ab values were on average higher than FO+IgG, there was no significant difference. On the other hand, strut spacing decreased significantly over time, shown by a reduction in signal in Fig. 5B, and there was a significant difference (Fig. 5D, p = 0.035), indicating that drug effects on mineralization can be captured within the void space of the pores.

The generation of the mask of the strut spacing and strut thickness also enables analysis of mineral density and content within the struts (Fig. 5E,F) and within the full volume of the model (Fig. 5G,H). The mineral density values represent the same metric as in Fig. 2D but without setting a threshold for mineral and restricting the analysis within the generated masks. The mineral content represents the cumulative sum of the density values multiplied with the corresponding volume of the mask. Across all metrics, treatment effects from DKK1Ab are observable as the average values of strut mineral density and content, as well as apparent mineral density and content of the DKK1Ab treated group were higher than in the IgG control, although a significant difference (p = 0.018) between the two groups was found only within the apparent mineral density (Fig. 5G) where the normalized value was 13.71% (p = 0.028) higher in FO + DKK1Ab than in FO + IgG at day 56.

### 2.5 Transcriptomic profiling reveals effects of Dickkopf-1 antibody at a molecular level in organotypic bone models

To gain insight into the molecular properties of the organotypic bone models, we performed RNA sequencing (RNA-seq) analysis. Principal component analysis (PCA) of the top 1000 most variable genes demonstrated a clear separation between sample groups, with the first principal component (PC1) explaining 86% of the variance and the second principal component (PC2) accounting for an additional 9% (Fig. 6A). This substantial separation along PC1 primarily distinguished samples based on patient origin (FO versus OI), while PC2 further separated samples based on treatment status. Notably, the gene expression profiles of DKK1Ab-treated and untreated OI organotypic bone models were more similar to each other than to those of FO models, regardless of treatment, indicating that patient-specific differences had a stronger impact on transcriptomic profiles than treatment effects.

**Fig. 6.**
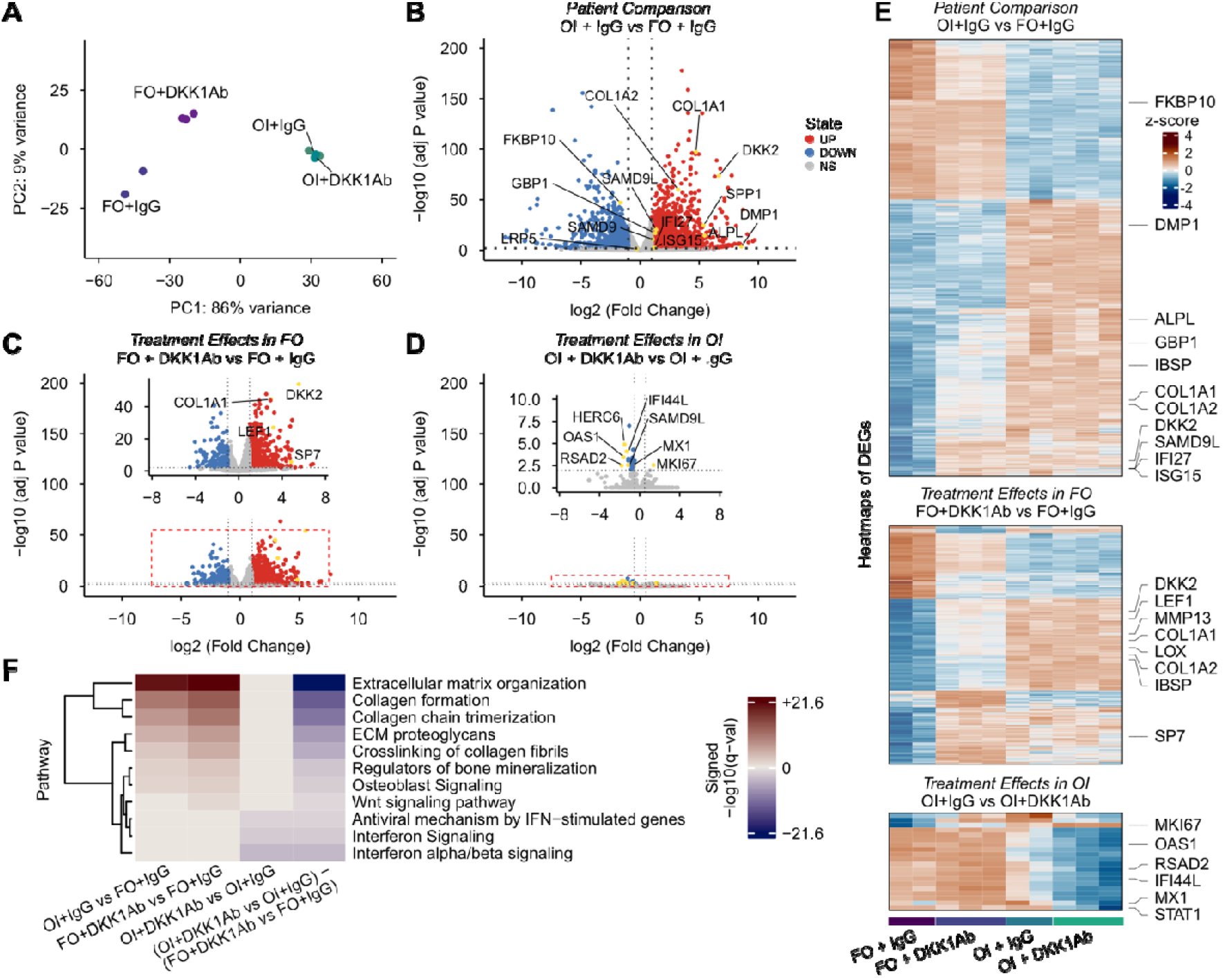
Endpoint transcriptomic analyses of treated organotypic bone models. (A) Principal component analysis of RNA-sequencing data based on the top 1,000 most variable genes across samples showing distinct clustering of samples by patient origin (FO versus OI) and treatment (DKK1Ab versus IgG control). PC1 (86% variance) primarily separates samples by patient origin, while PC2 (9% variance) further distinguishes treatment groups. Highlighted genes in volcano plots (B, C, D) represent key differentially expressed genes of biological significance. (B) Volcano plot showing differentially expressed genes (DEGs) between OI+IgG and FO+IgG control models, (C) DEGs in FO models following DKK1Ab treatment compared to IgG control. (D) DEGs in OI models following DKK1Ab treatment compared to IgG control organotypic bone samples identified by RNA-seq. FDR-adjusted P value < 0.01. (E) Heatmap illustration of gene expression values for DEGs in organotypic bone model groups. The colour gradient represents the z-score of log_2_ normalized counts, measured in counts per million (CPM). Red indicates a high positive z-score, indicating significantly higher expression levels; blue indicates a high negative z-score, indicating significantly lower expression levels. (F) Top upregulated pathways of all comparisons based on GAGE analysis.

Differential expression analysis revealed substantial differences between patient groups, with 2,522 differentially expressed genes (DEGs) meeting the threshold (|log2FC| > 1, FDR < 0.01), comprising 1,599 upregulated and 923 downregulated genes when comparing OI and FO control models (Fig. 6B, Supplementary Table ST-1).

Key bone-related genes, including DMP1, IBSP, DKK2, ALPL, and SPP1 were significantly upregulated in OI samples compared to FO controls, while the OI-causative gene FKBP10 showed differential expression, confirming the disease phenotype at the molecular level (Fig. 6B). Notably, OI organotypic bone models exhibited significant dysregulation of interferon-stimulated genes (ISGs) compared to FO controls. Among the upregulated genes in OI samples were key ISGs including SAMD9L, GBP1, ISG15, IFI27 and SAMD9, while other ISGs such as HERC5, TRIM5 were downregulated (Fig. 6B), indicating an interferon pathway-mediated inflammatory state in FKBP10-related OI.

Treatment with DKK1Ab elicited distinct transcriptional responses in FO and OI models. In FO models, DKK1Ab treatment induced significant changes in 746 genes (520 upregulated, 226 downregulated; Fig. 6C, Supplementary Table ST-2), with prominent involvement of Wnt signalling and bone-formation pathways. Among the most upregulated genes were COL1A1, COL1A2, DKK2, LEF1, SP7 and ALPL. In contrast, OI models exhibited only 20 differentially expressed genes (1 upregulated, 19 downregulated; (|log2FC| > 0.5, FDR < 0.01) Fig. 6D, Supplementary Table ST-3). The sole upregulated gene was MKI67, while the 19 downregulated genes were all interferon-stimulated genes (ISGs), including RSAD2, IFI44L, OAS1, and MX1.

The heatmaps of DEGs across individual samples for all three pairwise comparisons (Fig. 6E) revealed consistent clustering by both patient origin and treatment status. FO samples displayed uniform upregulation of osteogenic genes following DKK1Ab treatment, whereas OI samples showed limited transcriptomic shifts, reinforcing that treatment responses are pre-dominantly shaped by patient-specific background.

To interpret the biological relevance of these transcriptional changes, we performed Generally Applicable Gene-set Enrichment (GAGE) analysis using ConsensusPathDB. Across all four comparisons (Fig. 6F), we identified significant enrichment of pathways (q < 0.05), reflecting both disease-specific and treatment-specific transcriptional programs. The baseline comparison OI+IgG vs FO+IgG (Fig. 6F, Supplementary Table ST-4) showed that pathways for extracellular-matrix organization, collagen formation, and bone mineralization were enriched in OI. Treatment with DKK1Ab induced similar upregulation of these pathways in FO (Fig. 6F, Supplementary Table ST-5), while no significant upregulation of these pathways occurred in OI (Fig. 6F, OI+DKK1Ab vs OI+IgG, Supplementary Table ST-6). Instead, three interferon-related pathways were significantly downregulated in OI following DKK1Ab treatment. Finally, the interaction term (OI+DKK1Ab vs OI+IgG) - (FO+DKK1Ab vs FO+IgG) revealed an attenuated activation of extracellular-matrix, collagen-formation, and bone-mineralization pathways, together with an enhanced downregulation of interferon-related pathways in OI compared with FO (Fig. 6F, Supplementary Table ST-7).

### 2.6 Collagen expression, secretion, and deposition in organotypic bone models

To further investigate the effects of DKK1Ab treatment on bone matrix composition, we analysed collagen expression, secretion and deposition in both FO and OI organotypic bone models. Gene expression profiling revealed distinct patterns between FO and OI models, as well as treatment-dependent alterations (Fig. 7A). Notably, genes involved in collagen synthesis (*COL1A1, COL1A2, BMP1*) and mineralization (*DMP1, SPP1, BMP4*) were upregulated in both OI groups compared to FO IgG controls. Following DKK1Ab treatment, genes related to collagen synthesis were upregulated in FO organotypic bone models but remained unchanged in OI models, suggesting a differential regulatory effect.

**Fig. 7.**
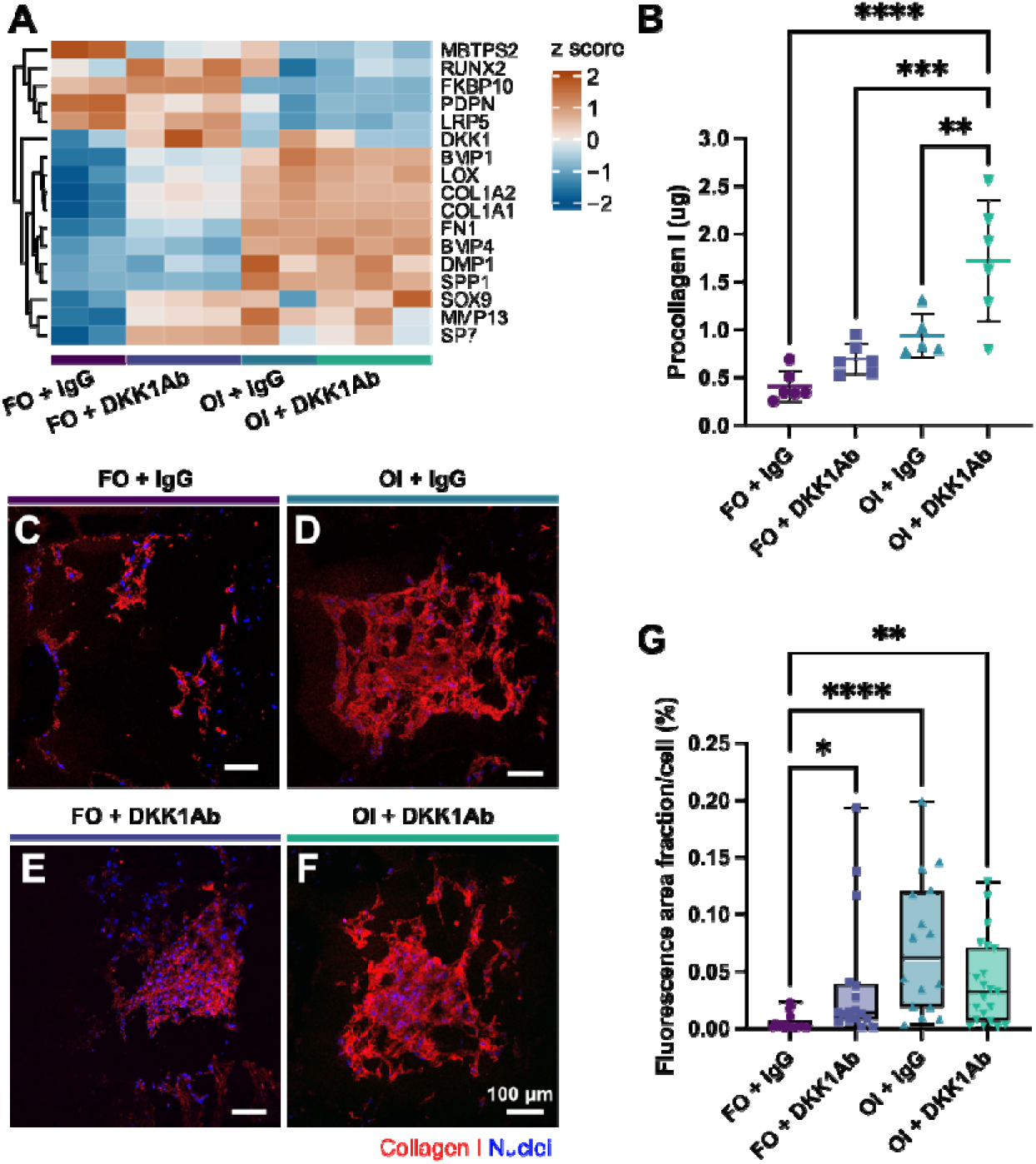
Collagen and mineralization-related gene expression and matrix deposition in organotypic bone models. (A) Heatmap depicting average gene expression related to collagen synthesis and mineralization in organotypic bone model groups: FO + IgG isotype control, FO + DKK1Ab treated, OI + IgG isotype control and OI + DKK1Ab treated. The colour gradient represents the z-score of log_2_ normalized counts, measured in counts per million (CPM). Brown indicates a high positive z-score, indicating significantly higher expression levels; blue indicates a high negative z-score, indicating significantly lower expression levels. (B) Intermediate quantification of procollagen I secretion measured by ELISA from cell culture medium taken 2 days after starting treatment. (C-F) Endpoint representative immunohistochemical images of collagen I (red) showing cell-secreted collagen in macroscale pores of cryosections of 3D bioprinted organotypic bone models. (C) FO + IgG control showing moderate collagen I staining with fibres organized along mineralized filament surfaces; (D) FO + DKK1Ab showing enhanced collagen I deposition; (E) OI + IgG control exhibiting dense collagen networks; (F) OI + DKK1Ab displaying similar collagen arrangement. Nuclei are counterstained in blue. Scale bar = 100 µm. (G) Quantification of collagen I fluorescence area fraction normalized to cell number. Data are shown as mean ± SD (B). Boxplots show median (center line), interquartile range (box), and min-max whiskers (G). Statistical significance between groups was identified by one-way ANOVA with Tukey’s multiple comparisons tests (B) or Kruskal-Wallis test followed by Dunn’s post hoc correction (G). *p<0.05, **p<0.01, ***p<0.001, ****p<0.0001.

These transcript-level changes were concordant with increased procollagen secretion (Fig. 7B) and collagen I deposition (Fig. 7C-G). Intermediate procollagen I levels were measured from cell culture medium collected two days post-treatment with 1 μg/mL DKK1Ab or 1 μg/mL IgG isotype control (Fig. 7B). OI organotypic bone models exhibited higher procollagen I levels compared to FO models. Treatment with DKK1Ab increased early procollagen I secretion, suggesting enhanced matrix production. Notably, OI + DKK1Ab models exhibited the highest procollagen I secretion among all groups (p < 0.0001 compared to FO + IgG), indicating that DKK1Ab treatment resulted in early enhanced collagen production in OI models. Collagen I deposition was also assessed at endpoint via immunohistochemical staining to visualize extracellular matrix organization (Fig. 7C-F) and corroborate transcriptomic findings at the protein level. FO control models showed moderate collagen I staining with fibres organized primarily along mineralized filament surfaces (Fig. 7C), while DKK1Ab treatment resulted in more intense and abundant collagen I deposition throughout the construct pores (Fig. 7D). OI models displayed dense collagen I networks in both treatment conditions (Fig. 7E, F). Quantification of collagen I fluorescence area fraction (Fig. 7G) confirmed significantly higher collagen signal in the pores of both treated and untreated OI models compared to FO + IgG. Following DKK1Ab treatment, collagen I deposition significantly increased in FO models, approaching levels seen in OI models.

Collectively, these results demonstrate that DKK1Ab treatment modulates gene expression and significantly increases collagen deposition in organotypic bone models, with effects more pronounced in the early secretion phase for OI models and in the accumulated matrix for FO models.

## 3 Discussion

This study demonstrates that mechanically stimulated 3D bioprinted organotypic bone models can reveal patient-specific therapeutic responses to DKK1Ab treatment in FKBP10-related osteogenesis imperfecta compared with metabolically healthy FO controls. Building on Akiva et al.’s pioneering work,^31^ our platform advances organotypic bone modeling by integrating mechanical stimulation^6,32,33^ and patient-specific disease recapitulation. This biomechanically dynamic environment uncovered phenotypes that remained invisible under static culture conditions, highlighting how mechanical forces shape cellular behavior and matrix organization.^34,35^

In our organotypic bone model we identified three distinct cellular microenvironments. Cells embedded within the mineralized hydrogel experienced direct cyclic compression and exhibited limited pericellular collagen, consistent with our previous observations in patient-derived osteoblasts.^32^ In contrast, osteoblasts seeded on the construct surface aligned along mineralized struts and produced organized collagen, parallel collagen fibers resembling lamellar bone. A third cell population spanning the construct pores formed dense, woven-like matrices, similar to those reports for hMSCs cultured on polymer scaffolds.^31,36,37^ This cellular heterogeneity was essential for assessing patient-specific *in vitro* responses to DKK1Ab.

Transcriptomic profiling was based on n=2 or n=3 samples per group. RNA sequencing revealed broad molecular differences between OI and FO constructs at baseline (2’522 DEGs at |log_2_FC|>1, FDR<0.01), consistent with prior reports of widespread transcriptomic remodeling in OI bone (approximately 3’700 genes)^38^. Whereas blood-based transcriptomic profiling of OI patients revealed an interferon-driven response characterized by roughly 100 DEGs^23^. Notably, DKK1Ab produced divergent effects: in FO, it upregulated classical osteogenic/Wnt targets including COL1A1, COL1A2, DKK2, LEF1, SP7 and ALPL^39^, while in OI, it upregulated MKI67 only, a canonical cell proliferation marker^40^, and downregulated 19 interferon-stimulated genes (e.g. RSAD2, IFI44L, OAS1, MX1). This selective transcriptional response suggests that for the FKBP10 OI patient, DKK1 blockade primarily mitigates inflammatory repression and enhances proliferative activity rather than initiating a full canonical Wnt program. The substantial overlap between gene sets distinguishing OI from FO constructs and those induced by DKK1Ab treatment in FO further suggests that Wnt-associated pathways regulating collagen synthesis are already upregulated at baseline in OI, limiting the additional transcriptional impact of DKK1 blockade. The result aligns with the work of Zhytnik et al. ^23^, who reported consistently elevated expression of IFI27, IFITM3, RSAD2, and GBP7 in COL1A1-OI patient blood.^23^ At baseline, the organotypic OI bone models showed a bidirectional interferon signature, with upregulation of SAMD9L, ISG15, IFI27 and GBP1 and downregulation of other ISGs. The direction of effect on osteoblasts differs between these genes. SAMD9 and SAMD9L act as growth suppressors, and variants in these genes have been associated with skeletal abnormalities,^41^ while GBP1 can limit cell proliferation and collagen synthesis^42^. ISG15 sustains osteoblast differentiation, since its knockdown reduces alkaline phosphatase activity, osteogenic gene expression and matrix mineralization in primary osteoblasts.^22^ The genes downregulated by DKK1Ab in OI map onto the same distinction. They include STAT1, which restrains osteoblast differentiation by sequestering RUNX2,^21^ and the growth suppressors SAMD9 and SAMD9L, whereas ISG15 was unchanged. The reduction of SAMD9 and SAMD9L is consistent with MKI67 being the only upregulated gene. The present analysis measured transcript abundance, while the functional evidence for ISG15 derives from murine osteoblasts^22^. Protein levels and functional consequences in the patient-derived constructs were not assessed.

OI models showed elevated COL1A1 and COL1A2 expression, accompanied by increased expression of collagen processing enzymes, including prolyl hydroxylases (P4HA1, P4HA2, P4HB), lysyl hydroxylases (PLOD1-3), and crosslinking enzymes (LOX, LOXL1-4). The co-ordinated upregulation of genes involved in collagen synthesis, processing, and mineralization is consistent with extensive remodeling of extracellular matrix production and maturation pathways in osteogenesis imperfecta, as previously described for altered collagen transport, post-translational modification, and biomineralization processes.^43,44^ FKBP10 function was impaired, consistent with a loss-of-function mutation. These molecular features were associated with elevated procollagen I levels in the cell culture medium and stronger collagen I signal within the extracellular matrix compared to FO controls. The accumulation of procollagen I in the culture medium is consistent with impaired collagen folding and crosslinking in FKBP10-related OI, which limits efficient incorporation into the extracellular matrix and promotes extracellular accumulation of immature collagen in cell culture media.^45^ Consistent with this, immunostaining of the bone explants (Fig. S3) revealed clear differences in collagen organization between OI and FO models.

The molecular response translated into distinct tissue phenotypes. Owing to the design of the 3D bioprinted scaffolds, the reseeded cells may experience macromolecular crowding within construct pores,^46^ trapping newly synthesized ECM within the pores. Cell distribution, morphology and, as a consequence thereof, ECM production^35^ was different between OI and FO constructs. Cells in FO constructs were found to line mineralized surfaces, adopting flattened morphologies mimicking lining cells, while cells in OI constructs spanned construct pores, adopting dendritic osteoblastic morphologies. This differential cellular behaviour resulted in surface cells producing an organized, layered matrix, while pore-spanning cells generated dense, woven-like matrices.^31^ OI constructs showed more pronounced pore-filling collagen by picrosirius red staining and increased collagen I immunostaining compared to controls. Overall, collagen differences may result from intrinsic patient-specific variations,^47–49^ local surface topography, ^50,51^ and morphology.^52^ These differences in collagen quantity and organization likely influenced subsequent mineralization patterns within the constructs.

At a tissue level, the organotypic OI bone models exhibited increased mineral density and altered mineralization kinetics compared to FO controls, consistent with the hypermineralization phenotype reported for OI bone.^53^ Time-lapse micro-CT also revealed a progressive increase in fracture score, notably only after reseeding, when osteoblasts actively deposited extracellular matrix under cyclic loading. The present study did not directly investigate the mechanisms underlying this structural deterioration. Mineralization patterns of reseeded constructs differed markedly, with homogeneous calcium deposition in FO pores versus large, distinct mineralized nodules in OI samples. Combined with upregulation of DMP1 (Dentin Matrix Acidic Phosphoprotein 1) and SPP1 (Osteopontin), these findings indicate dysregulated mineralization kinetics^54,55^, reflecting potential alterations in mineral crystal size and orientation^55^. The OI constructs also showed elevated baseline ISG15 expression alongside hypermineralization, compatible with the positive effect of ISG15 on osteoblast differentiation and matrix mineralization reported elsewhere.^22^ While the transcriptional response to DKK1Ab was limited, tissue-level micro-CT analysis showed lower median fracture scores in DKK1Ab-treated OI constructs than in IgG-treated OI constructs. However, the within-OI difference was not statistically significant. This pattern coincided with increased early procollagen I secretion, whereas endpoint collagen I deposition in OI constructs remained at the level of the IgG control.

DKK1 restricts Wnt signalling by binding the LRP5/6 co-receptor, and DKK1Ab is intended to relieve this inhibition.^16,56^ LRP5/6 gene expression was reduced in the tested OI organotypic models, in alignment with a documented LRP5/6 transport dysfunction in specific OI subtypes.^57^ Reduced availability of the co-receptor may therefore limit the response to DKK1Ab in these constructs. SP7/Osterix (OSX) showed a divergent response to treatment, which was increased in FO and decreased in OI.^58,59^ This inverse pattern is consistent with a disrupted OSX-DKK1 regulatory relationship, as previous studies demonstrate that OSX activates DKK1 expression and DKK1 expression decreases in the absence of OSX.^58,59^ LRP5/6 and SP7 were measured at transcript level only; direct measures of Wnt pathway activity were not included. Finally, our findings highlight the need to explore alternative therapeutic approaches for FKBP10-related OI, potentially targeting mechanisms beyond Wnt inhibition. Recent studies^60–62^ have shown that 4-Phenylbutyric acid (4-PBA) treatment reduces endo-plasmic reticulum stress and improves collagen folding in cells from OI patients with collagen type I mutations, suggesting its potential as a complementary approach for patients with protein folding defects.

Several limitations of the presented organotypic bone model should be acknowledged. With respect to biological complexity, the current system is a simplified monocellular model that does not yet incorporate the cellular complexity of bone such as osteoclastic cells or vascular components, both of which play essential roles in bone remodelling and nutrient transport in vivo, and endothelial and vascular smooth muscle cells have recently been identified as skeletal-disease-relevant cells that interact directly with the osteoblast lineage.^30^ Osteocyte maturation, including collagen-I maturation and lacunar-canalicular network formation, was established in this platform using human mesenchymal stem cells under long-term cyclic loading,^6^ but was not assessed in the present patient-derived osteoblast constructs. Because osteocytes are a principal source of Wnt antagonists in bone,^13^ it would be interesting to investigate in future studies whether DKK1Ab treatment influences osteocyte formation and dendritic network development in patient-derived constructs. Marrow-associated stromal elements are absent, and hierarchical matrix organization is reproduced only at the level of collagen fiber arrangement. Bone fragility and the response to osteoanabolic therapy in vivo depend on coordinated formation, remodelling, mineral maturation, and vascular support; the readouts reported here therefore characterize osteoblast-driven matrix formation and mineralization rather than integrated bone-organ physiology.

Reproducibility is influenced by batch-to-batch variation in bioink precursor materials, necessitating quality control and batch characterization to ensure consistency across experiments. Although the platform is not intended for high-throughput screening, its current scalability is limited by the need to load each construct in an individual bioreactor for 5 min three times per week. This is the most labor-intensive part of the workflow. Parallelization of cyclic loading across multiple constructs and transition to standard multiwell plate format compatible with mechanical stimulation would reduce manual handling and media requirements and support experimental cohorts comparable in scale to animal studies. At this scale, patient-derived constructs could generate structural, matrix, molecular, and transcriptomic endpoints within an eight-week workflow rather than the months often required for animal experiments.

In rare bone disorders, where donor numbers are inherently small, the intended application is reproducible assessment of treatment efficacy in cells from a single patient to support a personalized medicine approach rather than screening throughput. Validation across donors and interventions will nevertheless be required before the platform can be evaluated for patient-specific treatment assessment. In addition, RNA sequencing was based on expression profiles of n=2 or n=3 samples from donors of different sexes, offering exploratory views into transcriptomic differences between OI and FO samples. To elucidate functional implications of these gene expression alterations, results could be validated using RT-qPCR, ELISA, and immunostaining.^63^ Our findings in organotypic bone models highlight the central role of the extracellular matrix in tissue formation and maturation, consistent with broader evidence that matrix properties can direct cellular behavior.^64^ A thorough analysis is required to elucidate the signalling pathways governing our model’s development, which may in turn inform more effective therapeutic strategies for OI and related skeletal disorders.

Histopathological studies have shown that the proportion of woven versus lamellar bone varies with OI severity.^65^ Reflecting this complexity *in vitro* is challenging. In our study, the organotypic bone model employed cells from one patient and a single treatment concentration, limiting its ability to fully capture OI heterogeneity or predict treatment responses across broader patient populations. DKK1Ab was applied at a single concentration, and neither dose dependence nor target engagement was assessed. Future studies should include multiple patients with diverse OI mutations and explore different treatment concentrations and interventions.

The FO controls were not matched to the OI donor for age, sex, skeletal maturity, or clinical indication, because both cell sources derive from surgical waste material. Healthy donors of comparable age rarely undergo surgery, and when they do, it is most often following a major traumatic event. Consequently, the clinical circumstances leading to the availability of surgical waste tissue differ substantially between healthy controls and patients with OI, making meaningful matching for these characteristics impractical. Differences between FO and OI constructs can therefore not be attributed to disease status alone.

Developing effective treatments for rare diseases such as OI remains a significant challenge due to the complex pathophysiology, genetic heterogeneity, and limitations of *in vivo* disease models. Considering the diverse manifestations of OI bone, robust organotypic models are crucial for screening potential therapies and elucidating disease mechanisms in a controlled environment.^66,67^ Our model addresses this need by providing a patient-specific, mechanically stimulated platform that replicates key aspects of bone structure and function.

## 4 Conclusion

Our patient-derived organotypic bone model provides an experimental system that combines structural, matrix, molecular, and transcriptomic readouts from donor-derived bone-like tissue. OI constructs showed a bidirectional interferon signature, hypermineralization, and structural fragility. Following DKK1Ab perturbation, constructs from the tested OI donor showed a limited transcriptional response, downregulation of a defined subset of interferon-stimulated genes, increased MKI67 expression and early procollagen I secretion, and lower median fracture scores than IgG-treated OI constructs; the within-OI difference in fracture score was not statistically significant.

Beyond these findings, this platform integrates patient-derived biology with cellular- and tissue-level properties in a translational experimental format. It is intended for experimental cohorts comparable in scale to animal studies rather than for high-throughput screening. This reflects the setting of rare bone disorders, in which assessment of treatment efficacy in cells from individual patients is intended to support a personalized medicine approach rather than screening across large compound libraries. By linking patient-derived biology to multimodal readouts following a defined perturbation, the platform provides a translational framework for studying donor-specific features of FKBP10-related OI.

## 5 Experimental Section

### Patient-derived osteoblast isolation

Bone explants were collected from a 3-year-old female with type XI OI (FKBP10 c.890_897dup (p.Gly300Ter)) during an intermedullary rodding surgery and from a metabolically healthy 15-year-old male undergoing a femur osteotomy (FO) to correct limb malalignment. The bone explants were obtained from surgical waste material. All procedures were conducted in accordance with the Declaration of Helsinki. Written informed consent for the use of biological material for research purposes was obtained from the patient or from the patient’s legal guardians. Ethical approval was granted by Swiss Ethics (Kantonale Ethikkom-mission Zürich, KEK Nr. 2014-0300 and Nr. 2019-00811). Patient identities were protected, and all samples were fully anonymized by the children’s university hospital, with only sex, age, mutation, OI-type and site of collection made available for research. The bone explants were transferred from the operating room to the laboratory in sealed tubes containing Dulbecco’s Modified Eagle Medium (DMEM, Gibco) at room temperature and processed as described previously^32^ on the same day by rinsing in Phosphate Buffered Saline (PBS, Gibco), and vortexing three times in fresh PBS to remove blood. Directly, afterward osteoblasts were isolated based on tissue culture plastic adherence.^32^ Cells were expanded in DMEM supplemented with 10 % fetal bovine serum (FBS, Gibco), and 1 % antibiotic-antimycotic (Gibco, 100 U/ml penicillin, 100 mg/ml streptomycin, 0.25 mg/ml Amphotericin B), and cryopre-served in FBS and 10 % dimethyl sulfoxide (DMSO).

### Bone preparation

Segments of bone explants were also prepared for sectioning. After washing in PBS, segments of the bone explants were fixed on the same day in 4 % paraformaldehyde (PFA) for 48 hr, washed three times in PBS and decalcified in 12.5 % EDTA (pH 7.4–7.6) for 1-2 weeks at 4 °C (decalcification confirmed by scout view in micro-CT45 (SCANCO Medical AG, Brüttisellen, Switzerland)). Demineralized bone was then dehydrated in 20 % sucrose and 2 % Polyvinylpyrrolidone (PVP) for 3 days, followed by 30 % sucrose and 2 % PVP solution for 2 weeks. Bones were embedded in optimal cutting temperature compound (OCT, VWR) and flash frozen in liquid nitrogen. Cryosections (10–50 µm thickness) were prepared using Kawamoto’s cryofilm type 2C (SECTION-LAB Co. Ltd., Japan).^68^

### Bioink preparation and 3D bioprinting

Primary patient-derived osteoblasts were harvested and combined with hydrogels comprising 4.1 % (w/v) gelatine, 0.8 % (w/v) alginate, and 0.1 % (w/v) graphene oxide to form osteogenic bioinks with a concentration of 10x10^6^ cells/mL, as described previously.^24,32^ The bioinks were extrusion bioprinted using a 3DDiscovery bioprinter (RegenHU; Villaz-St-Pierre, Switzerland) equipped with a pneumatic dispenser. The lattice structures were printed directly onto double-sided tape (3M, Saint Paul, USA) secured on bioreactor platforms, following established procedures.^6^ Directly after bioprinting, samples were crosslinked in 2 % (w/v) calcium chloride in control medium (DMEM, 10 % FBS, 1 % antibiotic-antimycotic) for 10 minutes. Samples were washed twice in control medium and incubated (37 °C, 5 % CO_2_) overnight before being assembled into compression bioreactors.

### Organotypic bone intervention study

Metabolically healthy and *FKBP10*-OI (homozygous for FKBP10 c.890_897dup; p.Gly300Ter) cell-laden scaffolds were individually housed in custom bioreactors and subjected to uniaxial compressive loading three times per week in a mechanical stimulation unit (MSU). Each bioreactor was filled with 5 mL differentiation medium (DMEM, 10 % FBS, 1 % Anti-Anti, 50 μg/mL ascorbic acid, 10 mM β-glycerophosphate, 100 nM dexamethasone) and incubated at 37 °C and 5 % CO_2_ with media changes performed three times per week. Samples received fixed loading three times per week with a 0.07 N preload to define the sample contact position with the bioreactor piston, followed by a sinusoidal strain amplitude of 1 % at a frequency of 5 Hz for 300 seconds. Once per week medium was collected and bioreactors were scanned by micro-CT as described previously.^32^ After a preculture period of 4 weeks, 1 million fresh osteoblasts were pipetted onto each construct. After reseeding, samples received growth medium (DMEM, 10 % FBS, 1 % Anti-Anti, 50 μg/mL ascorbic acid, 10 mM β-glycerophosphate) supplemented with 1 μg/mL Dickkopf-1 antibody (treatment group) (R&D Systems AF1096) or 1 μg/mL Goat IgG (control group) (R&D Systems AB-108-C). Thereafter, to adapt to increased sample stiffness, the loading regimen was adjusted to an adaptive loading protocol (0.05 N preload, displacement-controlled position to 10 % strain, 1 % sinusoidal strain at 5 Hz for 300 seconds) for the final four weeks of culture.

### Time-lapsed micro-computed tomography

Mineralization was monitored by *in situ* time-lapsed micro-CT scanning as described previously. ^32^ Each bioreactor was scanned every 7 days in a micro-CT scanner (µCT45, SCANCO Medical AG, Brüttisellen, Switzerland) with a voxel resolution of 34.5 µm. The bioreactors were scanned at 45 kVp and 177 μA intensity with a 600 ms integration time.

### Image processing

Image processing was performed using an in-house software framework based on Python 3.12.11 (Python Software Foundation, Delaware). Time-lapsed micro-CT images were registered to the first time point for each sample using ANTsPy 0.6.1. A consistent volume of interest was established by defining a mask around samples. To diminish image noise, a constrained Gaussian filter was applied equivalent to IPL Scanco AG software V5.42 with sigma 1.2 and support 1. For segmentation of mineralized samples from the background, a global threshold of 74 mg HA/cm^3^ was applied. Mineral volume and density measurements were calculated from segmented greyscale images. Mineral formation rate was calculated by subtracting the mineral volume from the previous week. Samples that were broken were excluded from analysis. 3D visualizations were performed using ParaView 5.4.0 (Kitware, Clifton Park, New York).^69^

To quantify strut thickness and spacing of the bone model, we implemented a rigorous image processing pipeline similar to those described previously.^36,70^ The grayscale intensity images from the follow-up timepoints were registered to the images from day 28. These images were then Gaussian filtered (sigma = 1, support 1.2) to reduce noise and thresholded at 74 mg HA/cm³. Then an additional Gaussian filter with sigma = 3 and support 3 was applied using SciPy 1.15.0, followed by binarization with the threshold that was determined with Otsu (Scikit-Image 0.24.0) for day 28. This approach allowed us to define an envelope that tightly encloses the mineralized filament structure of the bone model.

To define a precise volume of interest, the binary masks underwent a series of morphological operations. First, holes within each XY plane were filled to ensure continuity of the structure. The masks were then dilated by 20 iterations to encompass nearby features, followed by an erosion of 21 iterations to create a tight-fitting boundary around the structure. Additional hole-filling was performed similarly in the other planes (XZ, YZ) but with each 4 iterations for dilation and erosion to generate a fully enclosed 3D envelope. This processing was performed for each time point individually, allowing consistent volumetric analysis across the experimental timeline. Morphological operations were implemented using NumPy 1.26.4 and SciPy’s 1.15.3 ndimage. The Boolean difference between this mask and the mask for the filament forms the mask for the strut spacing. Deformed and stable volumes were calculated by performing binary volume comparison on the time-lapsed image. A region that was stable across all time points was calculated from the intersection of the envelopes at each time point, combined with previously calculated masks for strut spacing and thickness. Strut thickness and spacing were calculated within the top 1 mm half of each sample according to the methods described by van Lenthe et al.^71^ with PoreSpy 2.4.3. The regional limitation to the top half was chosen to increase the sensitivity of the analysis to effects mediated by cells reseeded from the top.^36^ The strut mineral density was calculated by averaging the mineral density values of voxels within the region constrained by the mask for the strut thickness. The strut mineral content was calculated by calculating the cumulative sum of the product of density and volume for each voxel. The apparent mineral density and mineral content were calculated similarly but using the fully enclosed 3D envelope as a mask.

Fracture scores were calculated by combining volumetric deformity and changes in trabecular-like heterogeneity over time. A volume-based deformity index was derived from the ratio of deformed and lost volume to baseline volume, and cumulative heterogeneity change was quantified from percentile strut-spacing dynamics weighted by temporal rate. The normalized composite of these two terms yielded a unitless fracture score (0-100%), where higher values indicate greater structural compromise. A full mathematical formulation is provided in Supplementary Information.

### Scaffold mechanics

Scaffold mechanics were assessed using the in-house MSU following established procedures.^6,32,36^ Briefly, non-destructive measurements were acquired during the loading protocol. At endpoint, unconfined uniaxial compression tests were conducted under displacement control, applying a preload of 0.05 N to determine sample height and contact position, returning to baseline and then compressing with a displacement rate of 4 μm/s until the scaffold reached its yield point. During compression, both force and displacement were recorded and subsequently analysed using Python software (Python Software Foundation, Delaware, USA). The stiffness of the scaffold was derived from the fitted curve, calculated as the ratio of force to displacement at the steepest slope within the linear elastic range as described previously.^6^

### Sandwich ELISA assay

Procollagen Type I concentration was assessed in the cell culture supernatant using the Human Pro-Collagen I alpha I matched antibody pair kit (ab216064, Abcam) and ELISA accessory pack (ab210905, Abcam) according to the manufacturer’s instructions. The optical density was determined after 5 minutes of incubation with TMD Substrate (ab210902, Abcam) at 600 nm using a plate reader (Tecan Spark 10 M, Männedorf, Switzerland). Procollagen I concentrations were calculated from the collagen standard curves using GraphPad Prism 10.

### RNA Extraction, High Throughput Sequencing and Analysis

At endpoint, cell-laden scaffolds were washed thrice in PBS and removed from compression bioreactors. Constructs were homogenized in Trizol Reagent (Invitrogen, Thermo Fisher Scientific, USA) with a pellet pestle motor. Chloroform was added to induce phase separation and the clear aqueous phase was transferred to an RNeasy Micro Kit spin column (Qiagen, Switzerland) to isolate total RNA according to the manufacturer’s instructions. As an exploratory study, RNA sequencing was performed on n=2 IgG control samples and n=3 anti-DKK1 treated samples. RNA integrity was assessed using a 2100 BioAnalyzer from Agilent. Sequencing libraries were prepared with the TruSeq Stranded Total RNA Library Prep Kit from Illumina, following the manufacturer’s protocol. Paired-end sequencing was performed on all samples at the Functional Genomics Center in Zurich.

Raw sequence data quality was evaluated using FastQC (Version: FastQC v0.11.9, http://www.bioinformatics.babraham.ac.uk/projects/fastqc/). The raw reads were mapped to the human reference genome using subread (Version 2.0.1).^72^ The human genome index was created from subread-build index. The aligned reads were quantified on a gene basis using featureCounts (Version 2.0.1).^73^ RNA-sequencing data were uploaded to ArrayExpress (http://www.ebi.ac.uk/arrayexpress/) with the accession number GSE291327.

To explore the similarities and dissimilarities between the samples, the quantified count data were normalized using the Variance Stabilizing Transformation (VST) from the DESeq2 package.^73^ The principal component analysis (PCA) was performed on transformed read counts using the top 1000 most variable genes to assess the overall similarity between the sample. The differentially expressed genes analysis between FO + DKK1Ab versus FO + IgG, OI + DKK1Ab versus OI + IgG, OI versus FO + IgG groups was performed using DESeq2.^74^ The differentially expressed genes were selected using an FDR-adjusted P value cut-off < 0.01 and |log fold change| > 1, except for the comparison OI + DKK1Ab versus OI + IgG, where a relaxed cutoff (|log fold change| > 0.5) was applied due to the smaller dynamic range of response. The Ensembl ID was annotated to gene symbols and Entrez Gene using biomaRt (BioConductor version 3.20, R version 4.4.2) and only protein-coding genes, long noncoding transcripts (lincRNA antisense) and other processed transcripts with known function were retained. Lists of differentially regulated genes for the different group comparisons are provided in Supplementary Tables 1–3.

Gene-set enrichment analysis was performed using Generally Applicable Gene-set Enrichment (GAGE) (Bioconductor, version 3.20).^75^ For functional annotation of signalling pathways, we used gene sets from ConsensusPathDB.^76^ The analysis was performed on the basis of one-on-one comparisons, comparing between patients OI + IgG versus FO + IgG, and effect of treatment as FO + DKK1Ab versus FO + IgG and OI + DKK1Ab versus OI + IgG, data. The significant pathways terms were selected using an FDR-adjusted P value < 0.05. Lists of enriched pathways terms in upregulated gene sets are provided in Supplementary Tables 4-7.

### Sample preparation and histological staining

At endpoint, cell-laden scaffolds were washed twice in PBS and fixed in 4 % formaldehyde in PBS for 2 hours. Samples were washed twice and incubated in 10 % sucrose in PBS for 2 hours. Samples were incubated in 30 % sucrose in PBS overnight for cryoprotection. Subsequently, the scaffolds were embedded in optimal cutting temperature compound (OCT, VWR) and frozen in a methanol bath on dry ice. Samples were sectioned using Kawamoto’s cryofilm type 2C (SECTION-LAB Co. Ltd., Japan) on a cryotome (CryoStar NX70, Thermo Scientific).^68^ Prior to staining, 10-30 μm sections were affixed to microscope slides (SuperFrost™ Microscope Slides, ThermoScientific) using a solution of 1 % (w/v) chitosan in 1 % (v/v) acetic acid. Staining procedures included haematoxylin and eosin (H&E) staining to visualize cell nuclei, cytoplasm, and extracellular matrix, Alizarin Red S staining (2 mg/mL in acetone pH 4.3) (A5533, Sigma-Aldrich) to highlight mineralized extracellular matrix, and Picrosirius red staining (365548, P6744, Sigma-Aldrich) for collagen visualization. Imaging of the histological sections was conducted using an automated slide scanner (Panoramic 250 Flash II, 3Dhistech, Hungary) at ×20 magnification.

Collagen I protein was analysed by immunohistochemical staining. Briefly, 30 μm cryosections were washed thrice in PBS, followed by permeabilization in 0.1 % Triton-X-100 in PBS for 20 minutes. Subsequently, sections were washed again in PBS three times and blocked in 3 % BSA in PBS for 1 hour. Sections were incubated overnight at 4 °C with primary anti-collagen I antibody (1:200, ab6308, Abcam) in 1 % BSA in PBS. Control sections were incubated with mouse IgG isotype control (1:200, 026502, Invitrogen) overnight at 4 °C. Sections were washed thrice and incubated with secondary donkey anti-mouse Alexa Fluor 647 (ab150107, Abcam) for 1 hour. Sections were washed thrice in PBS and incubated with Alexa Fluor Plus 555 Phalloidin (1:500, A30106, Invitrogen) for 1 hour. Sections were washed thrice and incubated with Hoechst 33342 (1:200, B2261, Sigma Aldrich) for 20 min. Sections were washed thrice in PBS, mounted with ProLong Gold Antifade Mountant (P10144, Invitrogen), and sealed with nail polish. Imaging was performed using a Leica TCS SP8 confocal microscope (Leica Microsystems, Heerbrugg, Switzerland) and an automated slide scanner (Panoramic 250 Flash II, 3Dhistech, Hungary) at 40× magnification. Cell density (cells/mm^2^) was determined from Hoechst-stained nuclei automatically using ImageJ software. Similarly, fluorescence area fraction (%) in collagen I immunostained cryosections was evaluated using automatic Otsu thresholding in ImageJ.

### Statistical analysis

Statistical analysis was performed using GraphPad Prism 10 or R 4.2.2 with car, emmeans and nparLD packages. Data were first evaluated for normality (Shapiro-Wilk test) and homogeneity of variance (Levene’s test), followed by log_2_ transformation if assumptions for parametric testing were violated. *p-values less than 0.05 were considered statistically significant. Time-lapse data with complete repeated measures (no missing sample) were tested using repeated measures two-way ANOVA, followed by Tukey post-hoc correction when the group x day interaction was significant. Sphericity was evaluated using Mauchly’s test, and p-values were corrected using the Greenhouse-Geisser method when sphericity was violated. Single timepoint data were tested with one-way ANOVA followed by Tukey post-hoc correction. Non-parametric single time-point data were analysed with Kruskal-Wallis test followed by Dunn’s post-hoc correction. For longitudinal time-lapse data, treatment and time effects were assessed using rank-based analysis of longitudinal data (nparLD^77^) via the f1.ld.f1() function followed by Benjamini-Hochberg’s multiple comparisons adjustment.

## Supporting Information

**Fig. S1.**
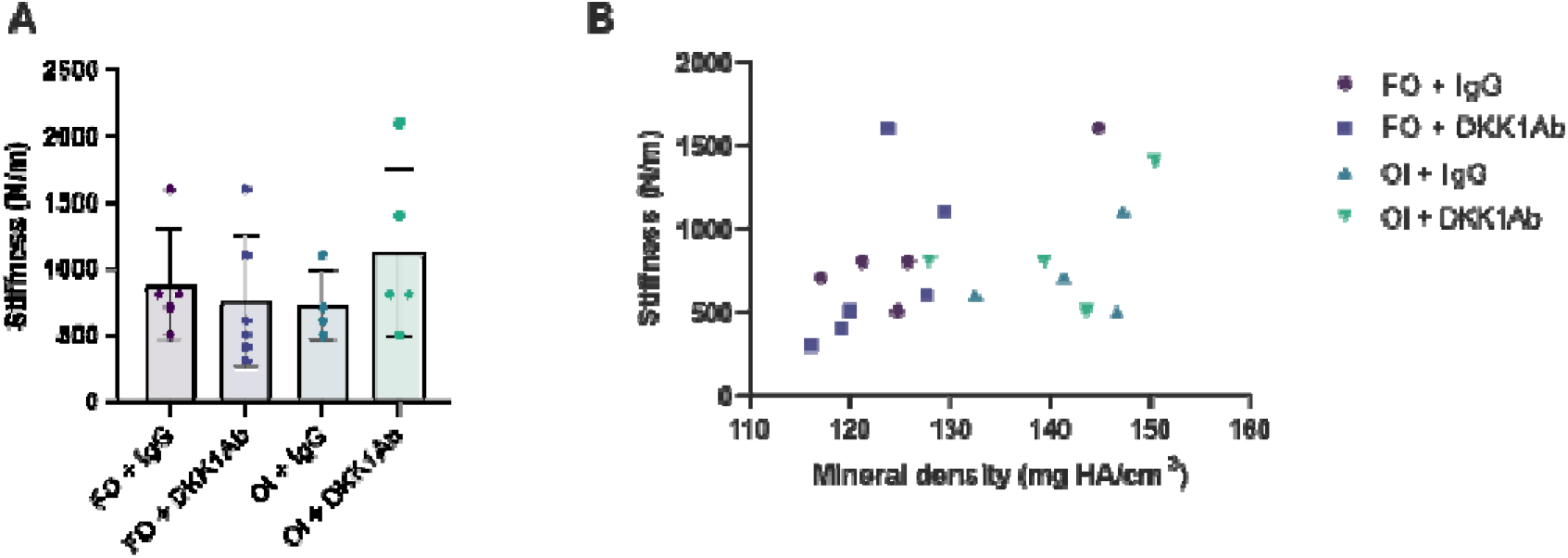
Endpoint mechanical assessment of organotypic bone models from a femoral osteotomy (FO) and an osteogenesis imperfecta (OI) patient. (A) Endpoint stiffness revealed no significant differences between groups (B) Correlation of stiffness to mineral density at day 56. FO samples show correlations similar to previous work,^6^ positive relationship between mineral density and resulting stiffness. OI sample mineral density is not correlated to stiffness.

**Fig. S2.**
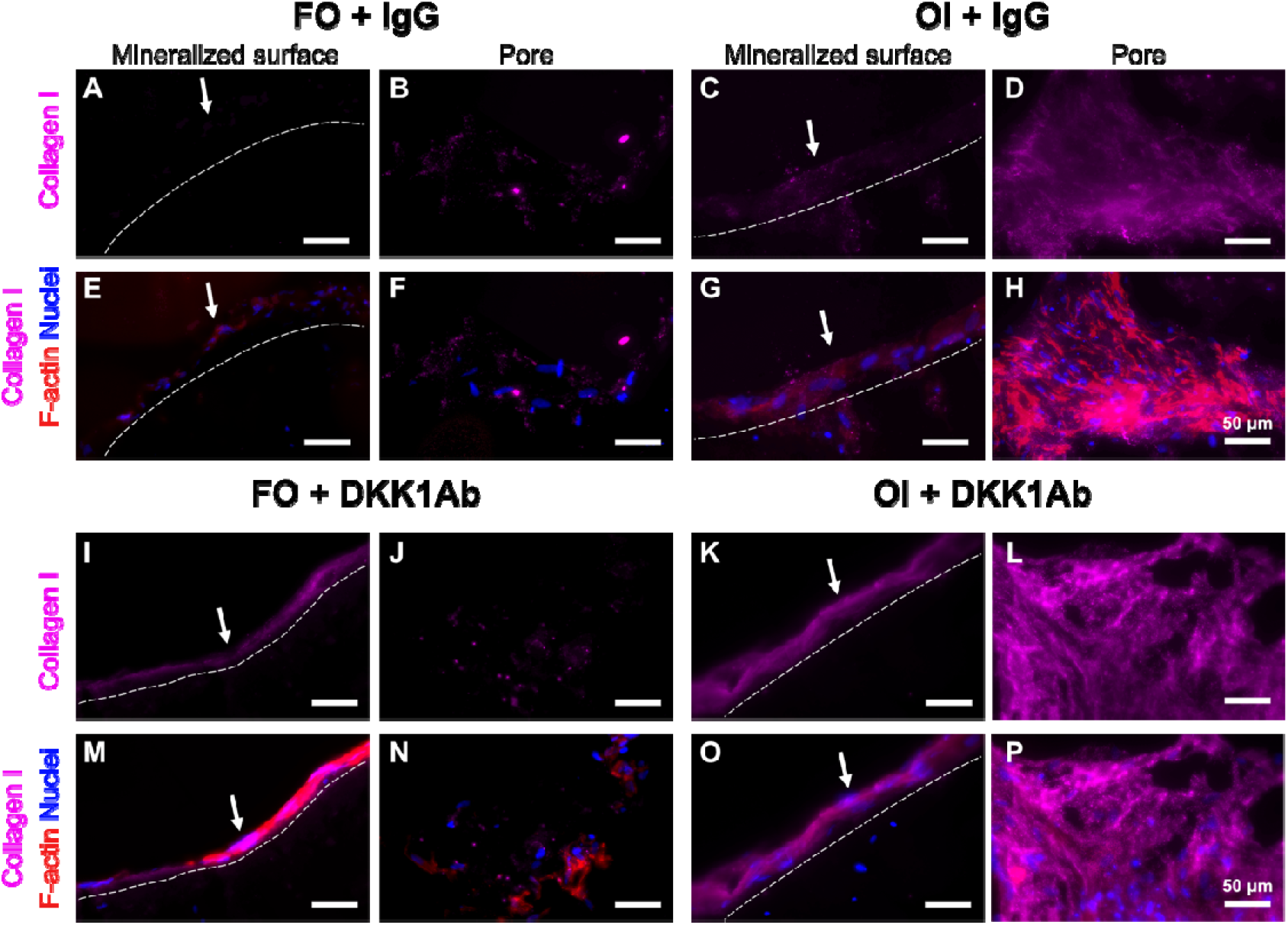
(A-P) Immunofluorescent staining of cell nuclei (blue), F-actin (red), and collagen I (pink) of sections of organotypic bone models from a patient undergoing femur osteotomy (FO) or from an osteogenesis imperfecta patient (OI) treated with IgG as control or DKK1Ab. Images show that cell-secreted collagen recapitulates the woven bone and surface modeling phenotypes (white arrows). The white dotted line indicates the mineralized surface. Scale bar = 50 µm.

**Fig. S3.**
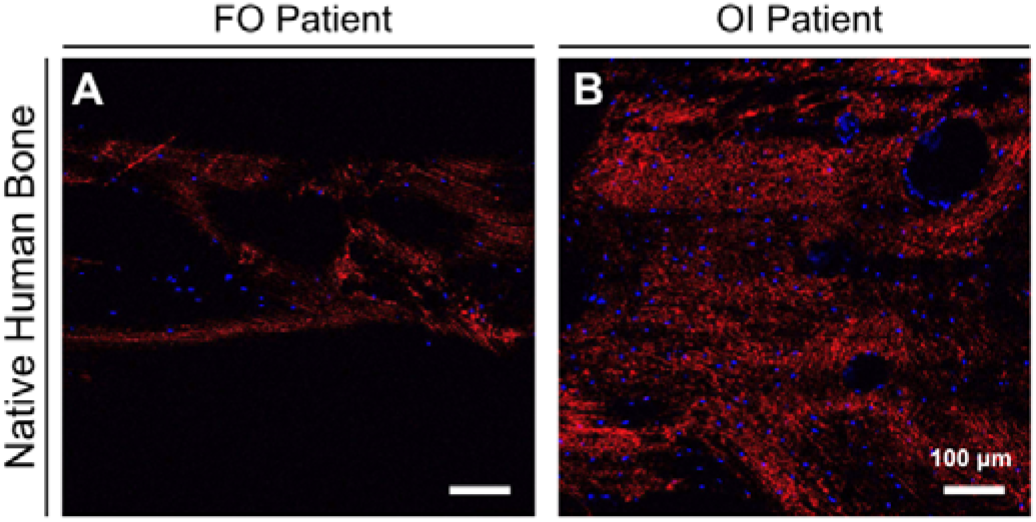
Confocal imaging of collagen type I in bone sections of FO and OI patients. A) Confocal imaging of immunofluorescent staining of cell nuclei (blue) and collagen I (red) of native bone from FO and B) OI patients. Samples serve as positive controls for collagen I immunostaining, revealing distinct differences in collagen organization and distribution in OI and FO bone. Scale bar = 100 µm.

**Supplementary Table S1.** (ST-1.csv): Differential gene expression (DEG) analysis of OI + IgG vs FO + IgG (FDR-adjusted p-value cutoff < 0.01)

**Supplementary Table S2.** (ST-2.csv): Differential gene expression (DEG) analysis of FO + DKK1Ab vs FO + IgG

**Supplementary Table S3.** (ST-3.csv): Differential gene expression (DEG) analysis of OI-DKK1 vs OI-IgG (FDR-adjusted p-value cutoff < 0.01)

**Supplementary Table S4.** (ST-4.csv): Gene-set enrichment analysis of OI+ IgG vs FO + IgG (FDR-adjusted p value cutoff < 0.05)

**Supplementary Table S5.** (ST-5.csv): Gene-set enrichment analysis of FO+DKK1Ab vs FO+IgG (FDR-adjusted p-value cutoff < 0.05)

**Supplementary Table S6.** (ST-6.csv): Gene-set enrichment analysis of OI+DKK1Ab vs OI+IgG (FDR-adjusted p-value cutoff < 0.05)

**Supplementary Table S7.** (ST-7.csv): Gene-set enrichment analysis of (OI+DKK1Ab vs OI+IgG)-(FO+DKK1Ab vs FO+IgG) (FDR-adjusted p-value cutoff < 0.05)

Part of the data discussed in this publication has been deposited in NCBI’s Gene Expression Omnibus and is accessible through GEO Series accession number GSE291327.

## Fracture Score

The fracture score is a unitless composite metric ranging from 0 to 100%, with higher values indicating greater structural compromise. The score integrates two primary components:

- Volume-based deformity index quantifying macroscopic structural deformation and loss
- Cumulative structural change measuring trabecular-like architectural changes.

The volume-based deformity index (*D_i_*) quantifies the percentage of baseline total volume that has become structurally compromised:

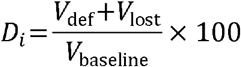

where *V*_def_ = deformed volume (mm³), *V*_lost_ = lost volume (mm³), *V*_baseline_ = baseline total volume at initial timepoint (mm³) and *D_i_*∊ [0,100] (%).

The cumulative structural change is calculated from a heterogeneity ratio *R*_90/10_ that describes the spread in strut spacing as:

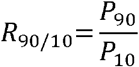

where *P*_90_ and *P*_10_ represent the 90th and 10th percentiles of strut spacing (μm). This ratio provides a robust measure of spatial heterogeneity, with *R*_90/10_ = 1 indicating uniform spacing and *R*_90/10_ > 1 indicating increasing heterogeneity.

For each sample j, the baseline heterogeneity ratio is defined as:

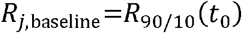

where *t_0_* represents the initial timepoint for sample *j*.

The absolute deviation from individual baseline at timepoint *t* is:

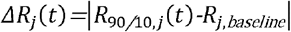

The rate of change in heterogeneity ratio between consecutive timepoints is:

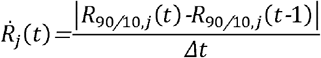

where *Δt* is the time interval between measurements.

A linear rate multiplier is applied to weight rapid structural changes:

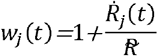

where R is the study-wide median rate of change for all positive rates.

The rate-weighted deviation at timepoint *t* is:

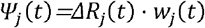

The cumulative structural change for sample *j* up to timepoint *t* is:

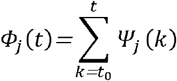

The cumulative structural change is normalized to a 0-100 scale using the maximum *Φ* within each sample

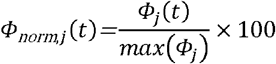

where *max*(*Φ*) represents the maximum cumulative structural change observed within each sample in the study.

The final fracture score integrates volumetric and structural components:

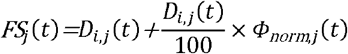

## Author contributions

Conceptualization: ADL, CS, GNS, PJL, MRo, CG, MRu, RM

Methodology: ADL, CS, GNS, AS, PJL

Investigation: ADL, CS, GNS, AS

Visualization: ADL, GNS, AS

Supervision: MRu, RM

Writing – original draft: ADL, GNS, CS, AS

Writing – review & editing: ADL, GNS, CS, AS, PJL, CG, MRu, RM

## Acknowledgements

This project was supported by the Strategic Focal Area ‘Personalized Health and Related Technologies’ (PHRT #2022-487 to GNS) of the ETH Domain (Swiss Federal Institutes of Technology) and the Swiss National Science Foundation (SNF Project 31003A_ 207542 to CG and MRo). The authors thank the Scientific Center for Optical and Electron Microscopy (ScopeM) of ETH Zurich for their support and assistance in this work. During the preparation of this manuscript the authors used the OpenAI ChatGPT and Anthropic Claude tool to improve the readability and grammar of the manuscript.

## Competing interests

GNS is an employee and shareholder of CompagOs AG. CS and RM are shareholders of CompagOs AG. All other authors declare that they have no competing interests.

## Notes

https://www.ncbi.nlm.nih.gov/geo/query/acc.cgi?acc=GSE291327

### Summary of Updates

We revised the manuscript to convey more clearly the focus and scope of the manuscript within health

